# A cosine similarity-based method to infer variability of chromatin accessibility at the single-cell level

**DOI:** 10.1101/315028

**Authors:** Stanley Cai, Georgios K. Georgakilas, John L. Johnson, Golnaz Vahedi

## Abstract

Cellular identity between generations of developing cells is propagated through the epigenome particularly via the accessible parts of the chromatin. It is now possible to measure chromatin accessibility at single-cell resolution using single-cell assay for transposase accessible chromatin (scATAC-seq), which can reveal the regulatory variation behind the phenotypic variation. However, single-cell chromatin accessibility data are sparse, binary, and high dimensional, leading to unique computational challenges. To overcome these difficulties, we developed PRISM a computational workflow and R package (https://github.com/stanleycai123/PRISM) that quantifies cell-to-cell chromatin accessibility variation while controlling for technical biases. Using data generated in our lab or publically available, we show that PRISM outperforms an existing algorithm, which relies on the aggregate of signal across a set of genomic regions. PRISM shows robustness to noise in low accessibility cells and reveals previously masked accessibility variation where accessible sites differ between cells but total number of accessible sites is constant. We also show that PRISM, but not an existing algorithm, finds suppressed heterogeneity of accessibility at CTCF binding sites. PRISM is a novel multidimensional scaling-based method using angular cosine distance metrics coupled with distance from the spatial centroid. PRISM takes differences in accessibility at each genomic region between single cells into account. This updated approach uncovers new biological results with profound implications on the cellular heterogeneity of chromatin architecture.

## 1 Introduction

Cell differentiation entails early lineage choices leading to the activation, and the subsequent maintenance, of the transcriptional program characteristic of each cell type (Natoli, 2010). The emerging theme from recent epigenomic studies is that alternative lineage choices involve the establishment of distinct accessible and open chromatin regions, a process instructed by lineage-determining transcription factors (Heinz et al., 2015). However, these findings rely on assays measuring an average of the chromatin states in thousands or millions of cells. Due to the binary nature of a regulatory element being open or closed in an individual cell, these bulk measurements further underscore the variability of chromatin accessibility at the single-cell level even in purified cell populations. Indeed, browsing the genome through the lens of bulk chromatin accessibility maps reveals large differences in the degree of openness across the genome, highlighting the need to study chromatin accessibility at the single-cell level.

Inference of cell-to-cell variability in the accessibility of regulatory elements is now possible with techniques such as single-cell assay for transposase accessible chromatin (scATAC-seq) (Buenrostro et al., 2015; Cusanovich et al., 2015; Jin et al., 2015). The signal measured by these assays at any genomic locus is fundamentally limited by DNA copy number and only 0, 1, or 2 reads can be generated from elements within a diploid genome. Hence, scarcity is intrinsic to these types of data. The exceedingly sparse nature of scATAC-seq data hinders studying the role of transcription factors in establishing the chromatin accessibly landscape at the single cell level. Thus, new computational methods incorporating the scarcity of data and the assay’s inherent bias are required. The current leading method for measuring cell-to-cell variation in chromatin accessibility, chromVAR (Schep et al., 2017), measures total accessibility in an individual cell at a set of DNA sequences unified by a common feature—such as binding events of a transcription factor. It then measures how much total accessibility differs from what is expected by calculating a technical bias-corrected Z-score for each cell. The standard deviation of these Z-scores constitutes the cell-to-cell variation in chromatin accessibility. However, an ensemble of cells can have similar total accessibility (*i.e.* number of accessible sites in a cell) yet be accessible at completely different regulatory elements. Thus, chromVAR is poorly equipped to handle chromatin accessibility variation in many cases as stated in the original paper (Schep et al., 2017).

We present PRISM, an R package for calculating cell-to-cell variation in chromatin accessibility using cosine similarity. Instead of measuring variation in total accessibility between two cells, PRISM measures whether two cells are accessible at the same set of regulatory elements using angular cosine distance. It then exploits principal coordinate analysis to measure how much each cell differs from the group norm for chromatin accessibility. Here, we demonstrate that PRISM outperforms chromVAR on various simulations when total accessibility is not varied, when signal is low, or when technical noise is high, in addition to real biological data publically available or generated in our lab. Together, PRISM can be used to construct a global and high-resolution view of epigenomic regulation in development and disease.

## 2 Methods

### 2.1 Chromatin accessibility variation prior to technical bias correction

We start by binarizing the accessibility at every site in each cell, such that 1 = accessible, 0 = inaccessible. Then we identify all sites unified by a common characteristic, such as TF-binding. Next, we plot every cell as a vector in space, with coordinates given by the binary accessibility at sites unified by this common characteristic. We next calculate how differences in chromatin accessibility between every pair of cells at these sites. Where *A* and *B* are binary accessibility vectors, the angular cosine distance is:

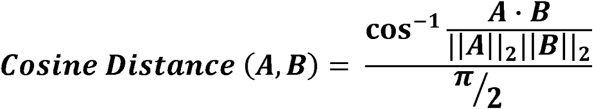

This can be seen as taking the angle between two vectors and dividing it by a normalizing factor of π/2.

Our next goal is to calculate how different every cell is from the group norm. We center the cosine distance matrix by subtracting column and row means and adding the overall mean. Then we spectrally decompose the centered matrix to define principal coordinates, and map all accessibility vectors to full principal coordinate space. We identify the centroid of the accessibility vectors. Then, we calculate each vector’s distance from the centroid. Finally, we calculate the average distance from the accessibility vectors to the centroid using R package vegan (Oksanen et al., 2017).

### 2.2 Correction for technical biases

Variation associated with technical factors such as GC content and mean accessibility differences can often introduce obstacles in interpreting NGS data. To overcome such limitations, for every original peak, we selected 30 “background peaks.” The set of peaks not bound by any TF were divided into 2-percentiles based on GC content. Every original peak was subsequently placed into a 2-percentile, and 30 background peaks within a 2- percentile of GC content were randomly subsampled with replacement. All background peaks were also within +/- 0.01 of the overall mean accessibility of the original 30 peaks.

We controlled for technical biases as follows:

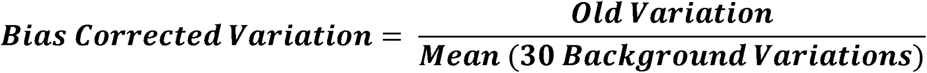

To measure accessibility variation beyond background noise, we calculated accessibility variation (with technical controls) for 30 randomly selected subsamples of peaks bound by no TF. Each subsample had the same number of peaks as the original peak set. This can be viewed as a negative control. Then we accounted for background noise as follows:

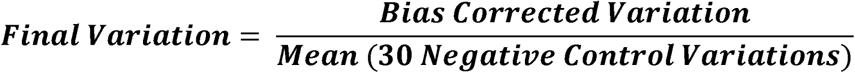

A variability equal to 1 implied that a TF was associated with no more variation than background noise. A variability below 1 implied that a TF was associated with less variation than background noise, and a variability above 1 implied greater variation than background noise. Additionally, the 30 Negative Control Variations were used to generate Z-scores and p-values for the observed variation.

### 2.3 Artificial variability simulations

We performed two sets of simulations on both human AML and mouse T-cell types. Simulation 1 varied accessible sites but not total accessibility between cells. Simulation 2 varied both accessible sites and total accessibility between cells. There were two subtypes for every simulation. Subtype A was run on the 50 highest accessibility cells, whereas subtype B was run on the 50 lowest accessibility cells.

In Simulation 1, 500 peaks were randomly sampled. Next, 500 GC-matched peaks were selected. To do this, the set of peaks excluding the original peaks were divided into 2- percentiles based on GC content. Every original peak was subsequently placed into a 2- percentile, and a GC-matched peak within a 2-percentile of GC content was randomly sampled.

Next, data for both peak sets were mixed. Accessibility matrices were artificially generated with peak data for the original 500 peaks replaced with peak data for the 500 GC-matched peaks in a proportion of the cells. Then variation (including background peak selection independent of the GC-matched peaks) was calculated for the artificially simulated accessibility matrix. The proportion of cells with original peak data varied from 100%, to 98%, to 96%, in 2-percentiles all the way down to 50% and 0%. By the Central Limit Theorem, in Simulation 1, the 2 peak sets do not differ significantly in total accessibility.

In Simulation 2, total accessibility was also varied. The same procedure as Simulation 1 was repeated, but the GC-matched peaks were drawn from peaks exclusively >75^th^ percentile in mean accessibility compared to other peaks.

### 2.4 K562 accessibility variation

Single-cell accessibility data and ChIP-seq data for 139 transcription factors were downloaded from ENCODE. PRISM and chromVAR were run on each transcription factor’s binding sites.

## 3 Results

### 3.1 The PRISM algorithm

We introduce PRISM for estimating cell-to-cell variation at the level of chromatin accessibility. Measurements of an ensemble of cells can show similar total level of accessibility within cells (*i.e.* comparable number of accessible sites in a cell) yet accessibility can occur at non-overlapping regulatory elements (Figure S1). Unlike chromVAR, PRISM infers cell-to-cell variability by capturing differences in accessibility of every genomic region of interest across individual cells. Our algorithm takes binarized read counts of individual cells and calculates variation of open chromatin across single cells at DNA sequences unified by an annotation such as transcription factor binding events (*i.e.* a set of peaks). PRISM then represents each cell in a high-dimensional space as a vector (Figure 1A). Each coordinate in the vector corresponds to accessibility of a regulatory element. To measure how different any two cells are at a given group of genomic regions, PRISM calculates the pair-wise angular cosine distance or the angle between two vectors (Figure 1B). While other distance metrics such as Euclidean distance can be used, it has been shown that angles are more robust in high-dimensional spaces (Li et al., 2015). The pair-wise differences between cells are then used to measure how different every cell is from the group “average”: Each cell is plotted as a point in principal coordinate space such that the Euclidean distance between two points (cells) is equal to the original angular cosine distance between two vectors. PRISM then finds the centroid of this unique point configuration. Every point’s distance from the centroid is calculated, and then these distances are averaged. This can be seen as each cell’s distance from the group norm for chromatin accessibility, and constitutes our measure of cell-to-cell variation prior to technical bias correction (Figure 1C). Our proposed method scales linearly with heterogeneity, in contrast to average angular cosine distance.

**Figure 1:**
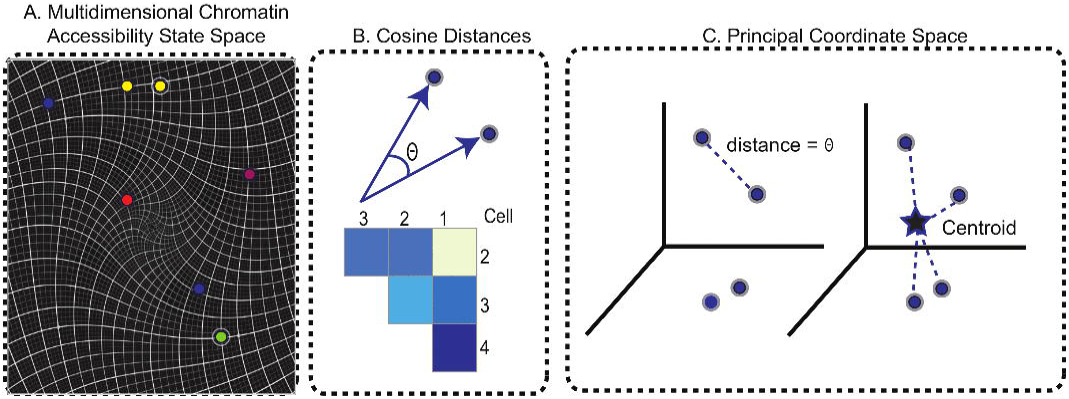
PRISM’s workflow for quantifying chromatin accessibility variation. (A) Each differentiating cell can be seen as a ball rolling down a hill, entering various furrows that represent different developmental trajectories. Here, each cell’s state is given by its chromatin accessibility across a transcription factor’s binding events. In mathematical terms, each cell’s coordinates are given by its chromatin accessibility levels at transcription factor binding events. (B) To measure how different any two cells are in terms of chromatin accessibility, we measure the angle between their vectors. A larger angle implies the two cells are more different in chromatin accessibility. As there are many possible pairs of cells, we form a matrix of cosine distances between cells. (C) To measure how variable the cells are as a whole with respect to accessible chromatin landscape, we perform principal coordinate analysis. Each cell is now plotted as a point in space. The Euclidean distance between any two cells is equal to the angle between their vectors. This specifies a unique point (cell) configuration in space and the centroid of the cells (points) is further calculated. Each cell’s distance from the centroid is measured. Then the average distance from each cell to the centroid constitutes the chromatin accessibility variation. Subsequent steps correct for technical biases.

To account for technical biases, 30 “background” sets of peaks are identified for every set of genomic regions. The background peak sets are matched for peak number, overall mean accessibility, and peak-for-peak GC content to the original peak set. Using the procedure outlined above, accessibility variation for each background peak set is calculated. The variations of the 30 background sets are then averaged. To obtain the bias-corrected variation, the variation of the original peak set is divided by the average variation of the background peak set with matching mean accessibility and GC content. After correcting for technical biases, a negative control is developed: 30 sets of peaks are randomly selected, each with equal peak number to the original peak set. The bias-corrected variation of each negative control peak set is calculated. Then the bias-corrected variation of the original peak set is divided by the average of the 30 negative control peak sets. This measures cell-to-cell variation in chromatin accessibility in units of background noise. A calculated variation equal to 1 implies that a chromatin feature is associated with equal variation to background noise. Together, unlike the previously proposed method chromVAR (Schep et al., 2017) which relies on differences in the aggregate of accessibility across a set of peaks between cells, PRISM takes differences in accessibility at each genomic region between single cells into account.

### 3.2 Generating synthetic heterogeneity in single cell chromatin accessibility and evaluating PRISM

To test whether our computational workflow could estimate cell-to-cell variability of chromatin accessibility, we generated synthetic heterogeneity within single cell chromatin accessibility maps using 2 models. While model 1 synthesizes heterogeneity assuming comparable levels of total accessibility between individual cells at a set of genomic regions, model 2 captures cell populations with large differences in total accessibility among cells. As genomic regions of interest, we randomly selected 500 peaks from ~50,000 open chromatin regions in human AML cells (Corces et al., 2016) and in mouse T cells. Relying on the central limit theorem, the randomly selected original and GC-matched peaks in model 1 comprise comparable levels of average accessibility across individual cells. The synthetic heterogeneity is then created by titrating the number of cells with the original or GC-matched peaks. This approach leads to a mixture of cells containing varying percentages of original versus GC-matched peaks (Figure 2A). The rationale to mix peaks rather than cells of different types is to prevent confounding factors such as differences in cell lysis affecting our assessment of cell-to-cell variability. We expect that when cells contain only original (or GC-matched peaks), the variability should be at a minimum. In contrast, when roughly half of cells are original peaks, the variability should be at a maximum. Based on how mixing of cells is controlled, we expect an inverse-U or concave shape for our measure of variability, peaking at around 50-50 original peaks and GC-matched peaks. We further synthesize heterogeneous data using model 2 which relies on the same procedure as model 1, except that GC-matched peaks are drawn from peaks with greater than 75^th^ percentile in mean accessibility compared to all other peaks. In other words, model 2 assumes the presence of a significant difference in total accessibility between cells underscoring a larger degree of cellular heterogeneity. We further augmented each model to have subtypes A and B such that subtype A was run on cells with highest accessibility in contrast with subtype B on lowest accessible cells. While subtype A is the most robust measurement and reflects an ideal sequencing coverage, subtype B tests the method’s sensitivity to technical noise. Together, model 1 is built such that heterogeneity is not caused by differences in total accessibility between cells, simulating cases where an ensemble of cells can have similar total accessibility but accessibility can occur at completely different regulatory elements (Figure S1). We further propose model 2 which captures cases where a major difference exists in the total accessibility of cells at genomic regions of interest (Figure S1).

**Figure 2:**
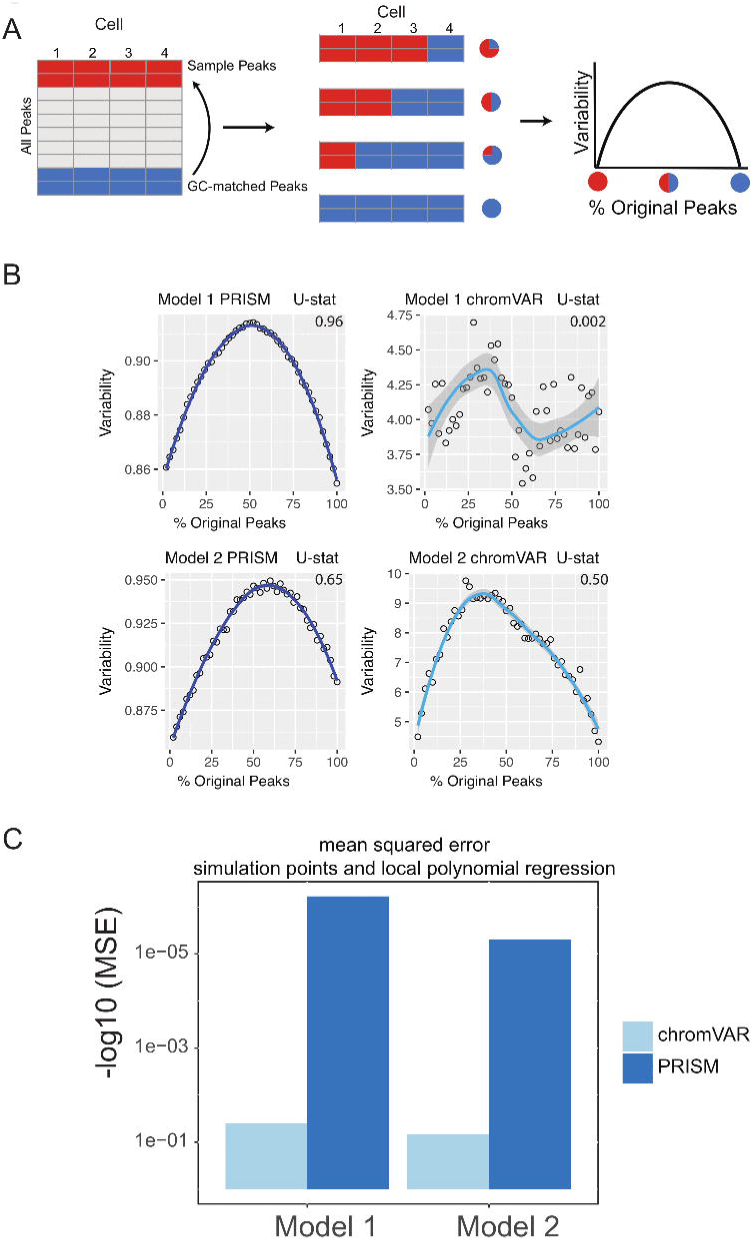
Simulations of cell-to-cell heterogeneity to compare PRISM and chromVAR. (A) We generated synthetic heterogeneity within single cell chromatin accessibility maps using 2 models. While model 1 synthesizes heterogeneity assuming comparable levels of total accessibility between individual cells at a set of genomic regions, model 2 captures cell populations with large differences in total accessibility among cells. As genomic regions of interest, we randomly selected 500 peaks from ~50,000 open chromatin regions in human AML cells (Corces et al., 2016) and in mouse T cells. Relying on the central limit theorem, the randomly selected original and GC-matched peaks in model 1 comprise comparable levels of average accessibility across individual cells. The synthetic heterogeneity is then created by titrating the number of cells with the original or GC-matched peaks. This approach leads to a mixture of cells containing varying percentages of original versus GC-matched peaks (Figure 2A). The rationale to mix peaks rather than cells of different types is to prevent confounding factors such as differences in cell lysis affecting our assessment of cell-to-cell variability. We expect variability to maximize when the data is a roughly 50-50 mixture of original and GC-matched peaks, and to minimize when data is completely original or GC-matched peaks, forming an inverse-U (concave down) shape. While subtype A is the most robust measurement and reflects an ideal sequencing coverage, subtype B tests the method’s sensitivity to technical noise. (B) PRISM outperforms chromVAR in two models. In model 1 subtype A, chromVAR does not conform to an inverse-U shape while PRISM does. In model 2 subtype A, chromVAR deviates from the curve of best fit more than PRISM. In order to see how well a simulation fit an inverse-U shape (concave curve), a test of concavity (U-statistic) was designed. The difference between variability of successive proportions of cells expressing original peaks was calculated. Then the Spearman correlation of this ordering with the decreasing number sequence 49 through 1 was calculated. This can be seen as checking to see if the derivative (slope) is continuously decreasing. Values close to 1 are ideal. (C) PRISM’s measurements were also significantly less noisy (stochastic) compared to chromVAR. To measure noise, we calculated the mean squared error (MSE), or average squared distance of each point from the LOESS curve. PRISM showed orders of magnitude smaller MSE values. The MSE is plotted on a square root scale.

We generated heterogeneous data using the two described models and calculated variability of chromatin accessibility at the single-cell level across the synthetic sets of peaks. We found an inverse-U shape in variability for the two models when variability was calculated using PRISM (Figure 2B). In order to see how well a simulation result fits an inverse-U shape (concave curve), a test of concavity was designed, which we termed the U-statistic. The difference between variability of successive proportions of cells containing the original peaks was calculated. Then the Spearman correlation of this ordering with the decreasing numbers 49 through 1 was calculated. This procedure checks if the derivative (slope) is continuously decreasing. Test of concavity (U-statistic) values close to 1 are ideal. We also measured each algorithm’s mean squared error (MSE) from its local polynomial regression (LOESS) curve. This assessed the degree to which an algorithm was susceptible to random fluctuations or noise. We further compared the values of variability across the simulated heterogeneous data as measured by chromVAR (Schep et al., 2017). We found that chromVAR falters under model 1 in the case that total accessibility is comparable across cells with a very low test of concavity U = 0.0002. In model 2, PRISM (U = 0.65) also conforms better to an inverse-U curve than chromVAR (U = 0.50). Notably, PRISM is significantly less noisy, with a mean-square-error (MSE) between the fitte several orders of magnitude lower than chromVAR (4.95*10^−6^ versus 0.07) (Figure 2C). PRISM further outperforms chromVAR in highly noisy conditions, such as in lowest accessibility cells (Figure S2A). These differences were reproduced when the synthetic heterogeneity was generated for scATAC-seq data generated in our lab in mouse primary T cells (Figure S2B). Altogether, testing multiple cases of controlled heterogeneity in publically available data in human AML cells and data generated in our lab in mouse T cells, we demonstrated that PRISM outperforms chromVAR in assessing cell-to-cell variability.

### 3.3 PRISM and chromVAR differ in predicting cell-to-cell variability in biological data

We next compared the predictions of PRISM and chromVAR on the effect of 139 transcription factors using real transcription factor binding data. Assessing cell-to-cell variability using the two methods revealed different predictions for 17 transcription factors in K562 cell line (Figure 3A). Among transcription factors that were reported differently between two methods, chromVAR but not PRISM inferred that CTCF binding events in K562 cell line could increase cell-to-cell variability at the chromatin accessibility level (Figure 3B). However, numerous studies mapping the genome-wide binding events of CTCF at the population level across a wide variety of tissues have shown the cell-type-invariant binding of this protein acting as an insulator, supporting PRISM’s prediction (Schmidt et al., 2012). On the other hand, several transcription factors including MCM family proteins were associated with high cell-to-cell variability by PRISM in contrast with chromVAR. Of note, the two methods were consistent in assessing variability at binding events of majority of transcription factors (Figure 3 C-D). Together, our simulation results titrating heterogeneity of chromatin accessibility at the single-cell level together with the application of biological data suggest that PRISM can infer cell-to-cell variability on the chromatin at the single-cell level and outperforms the existing method chromVAR.

**Figure 3:**
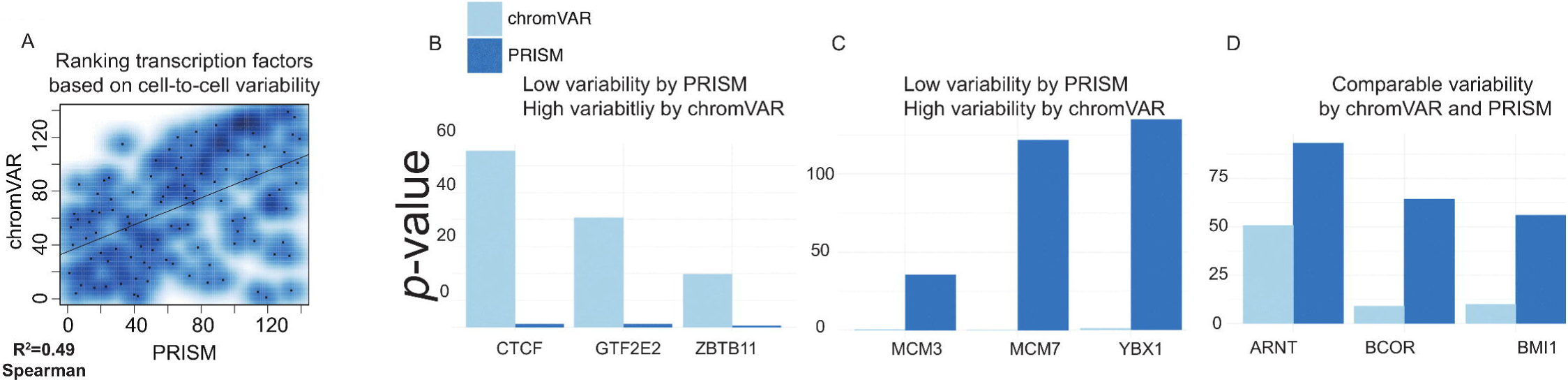
PRISM reveals previously masked chromatin accessibility variation on K562 cells. (A) Transcription factor binding data were extracted for K562 cell line using the Encode data. chromVar and PRISM are consistent in inferring cell-to-cell variability in 122 transcription factors while predictions are different for 17 transcription factors (R = 0.49). (B) We found six transcription factors with high rank of variability by chromVAR but low rank by PRISM. Three of these TFs including CTCF are shown. (C) Three transcription factors (MCM3, MCM7, and YBX1) were in the upper 25^th^ percentile in variability for PRISM, but found neutral by chromVAR. (D) PRISM and chromVar were consistent on the majority (122/139) of transcription factors examples include ARNT, BCOR, and BMI1.

## 4 Discussion

Existing functional genomic assays are inherently limited by the fact that they use bulk cells representing a weighted average of that population’s cellular constituents. Single-cell assays such as scATAC-seq help overcome several key obstacles that have frustrated efforts to assess the impact of transcription factors on chromatin at the single-cell level. Yet, these data present a number of intrinsic challenges, including systematic noise, scarcity and complexity of the data (Yuan et al., 2017). Recently, a method called chromVAR was developed to address these issues. chromVAR aggregates chromatin accessibility across peaks that share a common feature and assess the variability of the aggregate of accessibility between individual cells. While the aggregation of signal may address the variability of some single-cell data sets with certain statistical properties, this approach inherently masks heterogeneity within genomic regions across individual cells. To address this limitation, we developed PRISM a linear algebra-based method that takes into account the differences between every cell pair at individual genomic regions while correcting for GC-bias and average accessibility.

To evaluate the performance of PRISM and compare it with chromVAR in assessing variability of chromatin accessibility across single cells, we devised a computational experiment and shuffled real scATACseq data. Our framework generated synthetic scATAC data from the real measurements with various degrees of heterogeneity, which was further, used to evaluate the performance of PRISM. While PRISM predicted variability as heterogeneity increased, chromVAR failed to perform in cases where heterogeneity existed between peaks and across cells as a result of aggregating signal across all peaks (in particular model 1). We further showed that PRISM but not chromVAR can predict CTCF binding events to associate with low level of variability across individual cells. We propose that our methodology is able to effectively discover lineage-determining transcription factors establishing deterministic chromatin accessibility across cells of a lineage.

We have shown that our method, named PRISM, is able to overcome the obstacles in analyzing single-cell ATAC-seq, caused by the inherent nature of such assays, and provide a robust framework that assesses the effects of transcription factor binding on the chromatin accessibility, at the single-cell level. When compared to the state-of-the-art method, named chromVAR, we have shown that PRISM facilitated the discovery of lineage-determining transcription factors with the ability to preserve low variability of chromatin accessibility at the single-cell level.

---

**Figure S1:**
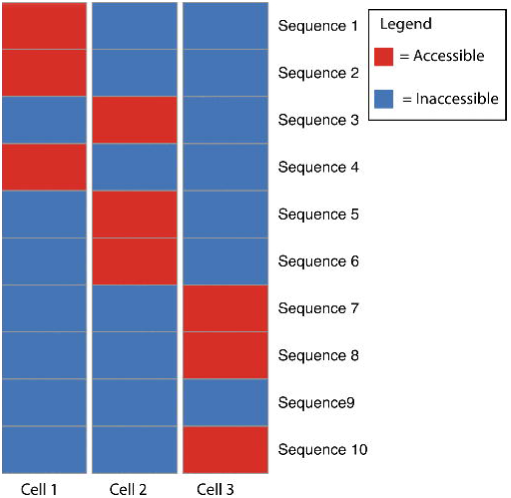
Chromatin accessibility variation can exist even if total accessibility is the same between cells. Measurements of an ensemble of cells can show similar total level of accessibility within cells (*i.e.* comparable number of accessible sites in a cell) yet accessibility can occur at non-overlapping regulatory elements. In this hypothetical case, red represents an accessible sequence in a given cell, and blue represents an inaccessible sequence. Each cell has 3 accessible sequences (red) total, or a total accessibility of 3. Thus, the total accessibility is the same between cells. But each cell is accessible at completely different DNA sequences, which may have different functions. Existing algorithms, such as chromVAR and Buenrostro *et.al* (2015)’s workflows, are built on the standard deviation of total accessibility, hence they cannot measure variation in these cases.

**Figure S2:**
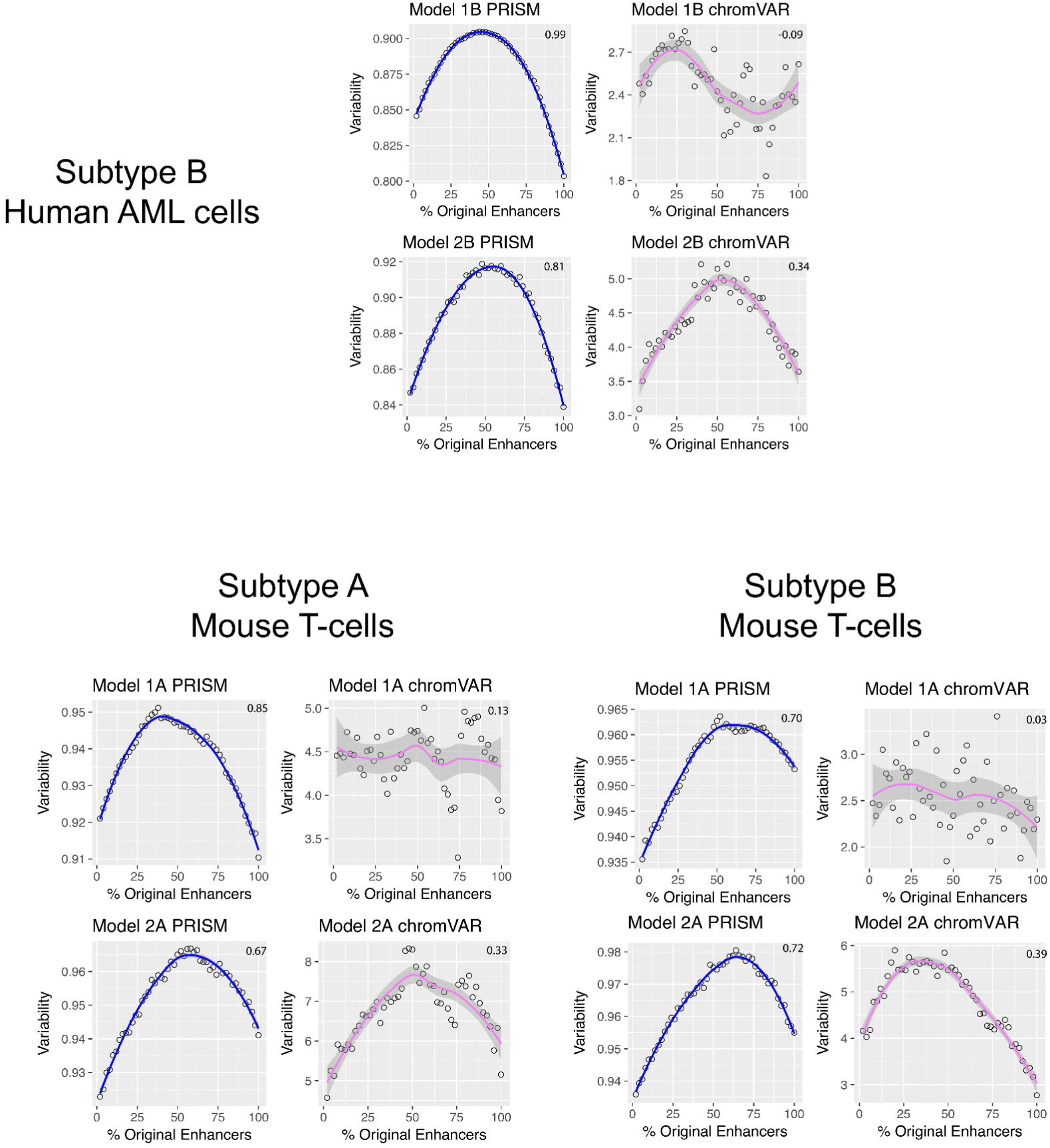
PRISM further outperforms chromVAR in various cell types and highly noisy conditions. (A) PRISM further outperforms chromVAR in highly noisy conditions, such as in lowest accessibility cells. While subtype A is the most robust measurement and reflects an ideal sequencing coverage, subtype B tests the method’s sensitivity to technical noise. (B) PRISM outperforms chromVAR in two models. In model 1 subtype A, chromVAR does not conform to an inverse-U shape while PRISM does. In model 2 subtype A, chromVAR deviates from the curve of best fit more than PRISM. In order to see how well a simulation fit an inverse-U shape (concave curve), a test of concavity (U-statistic) was designed. The difference between variability of successive proportions of cells expressing original peaks was calculated. Then the Spearman correlation of this ordering with the decreasing number sequence 49 through 1 was calculated. This can be seen as checking to see if the derivative (slope) is continuously decreasing. Values close to 1 are ideal. (C) PRISM’s measurements were also significantly less noisy (stochastic) compared to chromVAR. To measure noise, we calculated the mean squared error (MSE), or average squared distance of each point from the LOESS curve. PRISM showed orders of magnitude smaller MSE values. The MSE is plotted on a square root scale. (B) These results were reproduced when our synthetic heterogeneity was generated for data we generated in our lab in mouse primary T cells.

## Author Contributions

All authors contributed extensively to the work presented in this paper. GV conceived the project, administered the analyses and wrote the manuscript. SC developed and implemented the method. GG and JJ provided technical support and conceptual advice.

## Acknowledgments

The authors thank the sequencing facility at the Penn Vet Center for Host-Microbial Interactions. We thank members of the Vahedi and Faryabi labs particularly Dr. R. Babak Faryabi as well as Gregory Schwartz and Yeqiao Zhou for discussions.

## 1 Data Availability Statement

The datasets generated for this study can be found through NCBI GEO: GSE99159.

PRISM is an open source framework, freely accessible through Github (https://github.com/stanleycai123/PRISM).

